# Hybrid Chemomechanical Promotion of PEDOT Adhesion onto Flexible Microelectrode Arrays for Chronic Neural Stimulation

**DOI:** 10.1101/2020.09.09.289405

**Authors:** Mohammad Hossein Mazaheri Kouhani, Alexander Istomin, Proyag Datta, Neil H. Talbot

## Abstract

Advances in neural prosthetic technologies demand ever increasing novelty in material composition to enhance the mechanical and electrochemical properties of existing microelectrode arrays. Conductive polymers present advantages such as mechanical flexibility, outstanding biocompatibility, remarkable electrical properties and, most of all, cellular agreement. However, for long-term chronic applications, they fall short in their electrochemical endurance and mechanical adhesion to their substrate materials. Multiple electrochemical approaches have been investigated to improve the adherence of Poly(3,4-ethylenedioxythiophene) (PEDOT) to underlying metallic thin films. In this work, an electrochemical treatment of diazonium salt on platinum microelectrodes is incorporated as an electrochemical adhesion promoter for PEDOT and it is further combined with using the highly microporous geometry of Platinum Grey (Pt-Grey); a technology developed by Second Sight Medical Products Inc (SSMP). The intertwined mechanical integration of Pt-Grey and PEDOT molecules together with the covalent binding agency of diazonium salt demostrate a composite coating technology with long-term stability of more than 452 days while providing >70× enhancement to the interfacial capacitive impedance.

## INTRODUCTION

As incredibly complex as it may be to understand how the brain works, with collective cooperation and global collaboration, humans have uncovered remarkable insights into the molecular and electrochemical mechanisms that govern the central nervous system. The neuroscience and neuroprosthetic community demand an ever-increasing density of electrodes with more channels in the important mission of expanding the communication bandwidth between the two complex worlds of living neurons and electronic computers. Besides, simultaneous somatodendritic patch clamp recordings and two-photon imaging has recently shown that XOR operations are performed in human dendritic cells [1].

These findings are strong evidence that not only is it highly desirable to speak with and listen to each neuron separately with thousands of channels, but it can also be advantageous to tap into and modulate multiple points on a single neuron and treat them as more than singular units of summation and multiplexion. This possibility requires compact microelectrode arrays (<10μm) with minimal spacing (<5μm). This would allow placing thousands of individually controlled channels in a series of ultra-small penetrating microelectrode arrays with high aspect ratio providing ultra-high-bandwidth direct brain-computer-interface for neural modulation including both recording and stimulation.

The standard approach for increasing the density of electrodes is to decrease the size of individual electrode sites so that more can be placed on a given area of epicortical or penetrating electrode arrays. The challenge in doing so is to enhance the electrical charge storage capacity (CSC) of ultra-small electrode sites (<50um). Even though there are multiple approaches to enhance CSC of existing inorganic coating technologies [2–5], conductive polymers promise significantly better cellular agreement [6–8] while they maintain superior CSC, high flexibility, and lasting biocompatibility [9]. Among them, Poly3,4-ethylenedioxythiophene (PEDOT) stands out as the most established and investigated in literature to date. PEDOT’s advantages over solid state materials are in allowing the transportation of ionic charge across its entire bulk as well as the transport of electrons in an ohmic fashion.

These inherent advantages result in increased charge storage capacity, larger interfacial capacitance, and lower electrical resistance. PEDOT’s typical injectable charge into the surrounding tissue can be maximized to more than 15 mC/cm^2^ [10] which is approx. 100x better than platinum [11] and approx. 15x better than iridium oxide [12] and titanium nitride [13].

Another advantage of PEDOT in neural interface coatings is its customizability in accepting various electrical dopants or immunosuppressant drugs during its electrodeposition [14]. N. Kim et al. [15] showcase this advantage by studying and comparing more than ten secondary dopant molecules such as polystyrene sulfonate (PSS), perchlorate (CLO4), and hexafluorophosphate (PF6) integrated into PEDOT as counter ions under a diverse range of chemical reactions. While each of these molecules and reactions solve different problems and offer unique capabilities for diverse applications, in this work, tetrafluorborate (BF4) is incorporated as the secondary dopant material while electropolymerizing PEDOT.

As reported by Bodart *et al.* [16] PEDOT:BF4 is durable and offers favorable electrical properties; however, long-term and in-depth biocompatibility of PEDOT with any of these variations remain unknown. One of the main obstacles in adopting PEDOT for chronic neural stimulation is its instability and poor adhesion to inorganic electrode surfaces. Even though some researchers have reported attempts [17–18] to solve this problem and some others suggest reducing thin film shear stress helps in principle [19], the enhancement of long-term durability of these delicate thin films remains a challenge.

Diazonium salt has been shown capable [20] in establishing decently strong covalent bonds between the PEDOT thin films and their underlying metallic base material. In fact, this technique was initially tested using Scotch tape and it was observed that it was almost impossible to remove the PEDOT coatings on platinum when adhesion was promoted with diazonium salt. Whereas in the case of electrodeposition on smooth platinum without diazonium salt as an adhesion promoter, PEDOT thin films were easily peeled off from Pt sites with Scotch tape.

In this report, microporous Pt-Grey as an underlying thin film substrate is suggested and investigated for promoting the integration of PEDOT adhesion. Furthermore, diazonium salt is electrografted on Pt-Grey which yields a covalent bond between Pt-Grey and PEDOT on a molecular level [20]. Eventually, the unique hybrid advantages of PEDOT:Diaz:Pt-Grey composite is presented in attaining superior electrical characteristics while remaining unchanged for long periods of time under constant round-the-clock electrical stimulation *in vitro*.

This is the first time the mechanical integration of microporous material is combined with the electrochemical benefits of using diazonium salt in creating a hybrid composite that demonstrates long-standing strong adhesion between PEDOT and inorganic microelectrodes such as Pt. The major significance of this report is in highlighting the benefits of microporous solid-state coating technologies [21–27] as substrate materials for conductive material coatings while incorporating electrochemical adhesion promotion strategies such as diazonium salt.

## RESULTS

First, a polyimide-platinum microelectrode array was selected (Fig. 1-d) that had smooth platinum electrode sites (Fig. 1-e) which were coated with PEDOT:BF4 with and without diazonium salt for comparison. Next, a similar microelectrode array (Fig. 1-a) was picked that had Pt/Pt-Grey electrode sites (Fig. 1-f) which were coated with PEDOT:BF4 with and without diazonium salt for comparison. A representational figure showing a cross-section of the proposed hybrid composition is illustrated in Fig. 2-a which shows the mechanical integration of PEDOT coating into the maze that is created by the microporous geometry of Pt-Grey. Fig. 2-b shows the other case where PEDOT is directly deposited on smooth Pt electrode sites.

**Figure 1:**
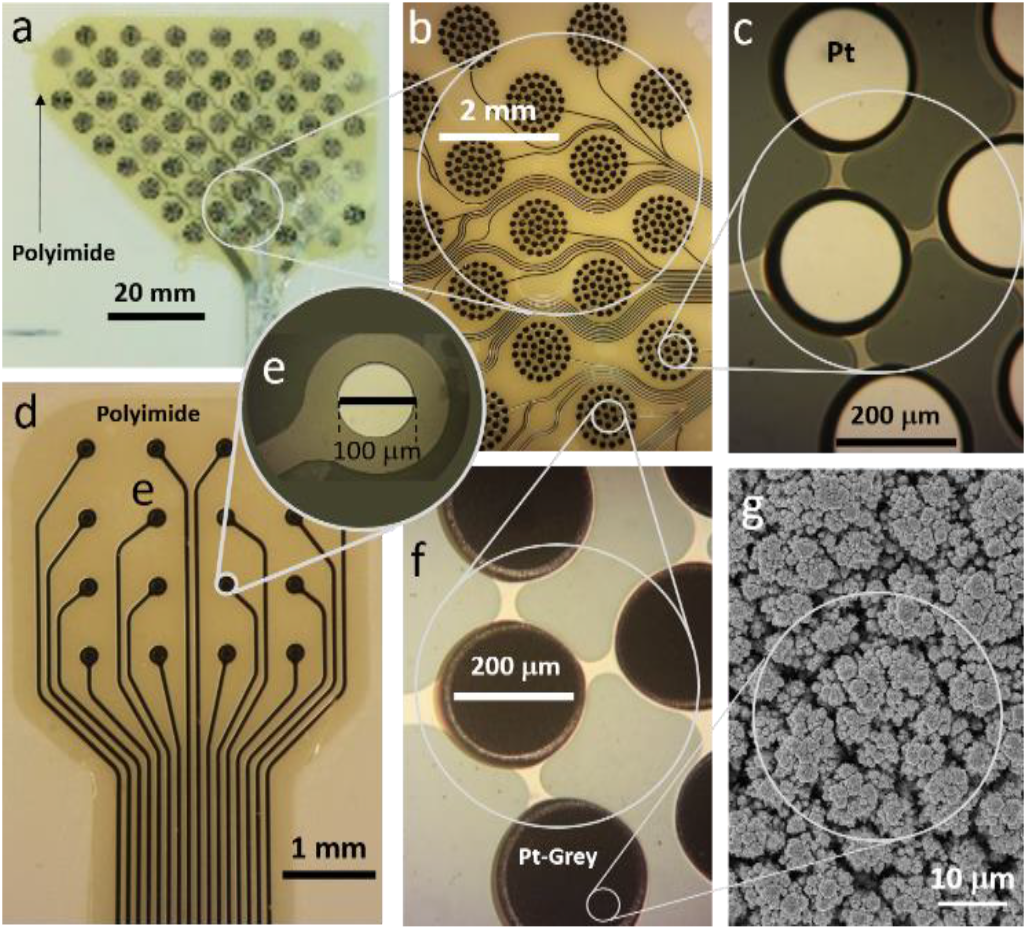
The epiretinal and epicortical electrode arrays used in the first and second experiments. a) The electrode array with 60 channels wired out for external connection. The array is sitting on a ceramic carrier for bench-top handling. b) A closer view of the same array under optical microscopy. Each cluster consists of 37 sites that are shorted together making up one channel. c) A closer view of the same array showing the circular platinum sites that are shorted together. d) The top view of a 16-electrode array. e) A closer view of one of the sites showing the Pt electrode at a diameter of 100 μm. f) A closer view of the 60 channel array showing the circular sites that are shorted together and coated with Pt-Grey. g) Scanning electron microscopy (SEM) image of Pt-Grey coating (adapted from D. Zhou *et al.* [28]).

**Figure 2:**
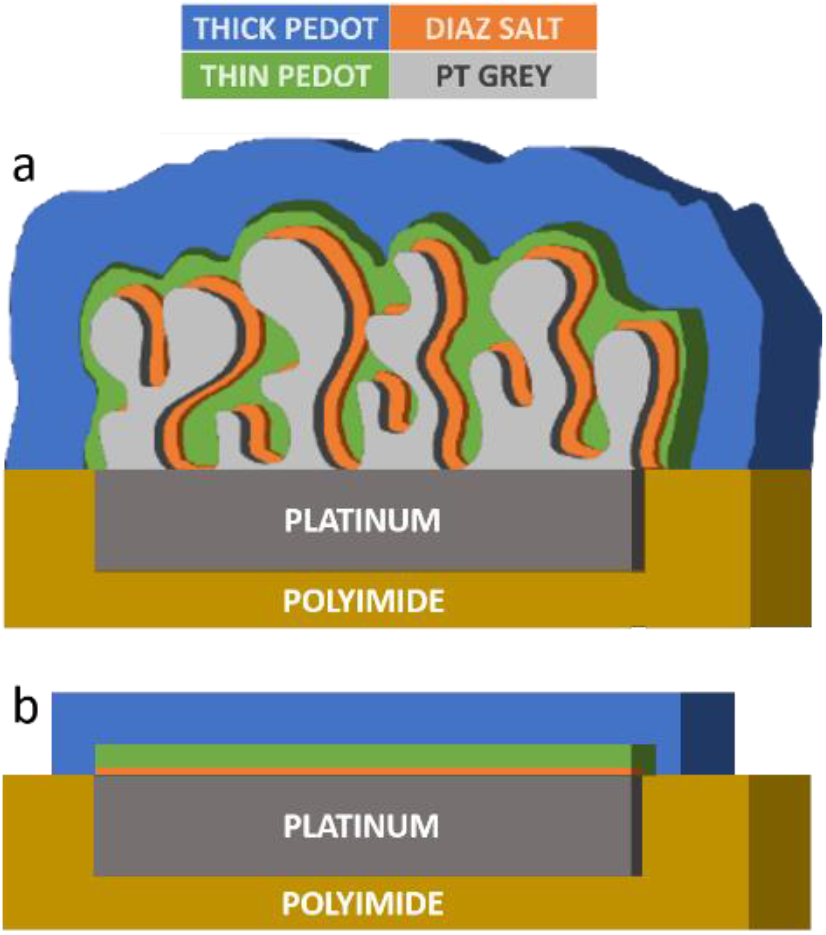
Schemcatic diagram illustrating the order of thick/thin PEDOT and diazonium salt on a) Pt-Grey coated on platinum electrode site and b) blank platinum electrode sites.

We galvanostatistically electroplated PEDOT on blank platinum electrodes (d=100μm) using different electrochemical currents to identify the optimum current density that creates smooth, thin and uniform PEDOT films. As shown in Table 1, increasing thicknesses of PEDOT are produced by increasing the applied DC current from 37.5nA to 62.5nA and 87.5nA for 2 minutes each. The values in Table 1 are averaged results from 4 electrode sites that were plated simultaneously in parallel. As shown in Table 1, while the improvement in the interfacial electrical resistance remains similar in all three cases, the electrochemical capacitance is enhanced seven-fold when the PEDOT is thickened three-fold. A diameter of 100μm for a circular/planar electrode site and the electrical current of 37.5nA yield optimum quality and correspond to a current density of 0.48 (mA/cm2) which was also employed in the next set of electroplating experiments. The cyclic voltammetry (CV) results of the platinum electrode sites before and after PEDOT coatings are illustrated in Fig. 3 at three different electrical currents with the same electroplating duration. The absolute area of the bottom half of the hysteresis loop in each case represents the amount of charge being stored inside the entire interfacial coating and transferred to its surrounding solution during each stimulation phase. Next, Pt-Grey electrode sites were electrografted by diazonium salt and then electroplating of various thicknesses of PEDOT was conducted on top. Table 2 shows the electrode sites with diameters of 200μm with and without Pt-Grey coating before and after coating PEDOT atop. From the first experiment the electrical current density of 0.48 (mA/cm^2^) was found to be optimal. In the second experiment (Table 2) the electrical current was kept constant at the optimum current density (0.48 mA/cm^2^) which corresponds to 150nA for the electrode sites with diameters of 200μm. Electoplating time durations of 120s, 140s, and 240s, resulted in capacitances of 18μC, 21μC, and 36μC, respectively.

**Table 1:**
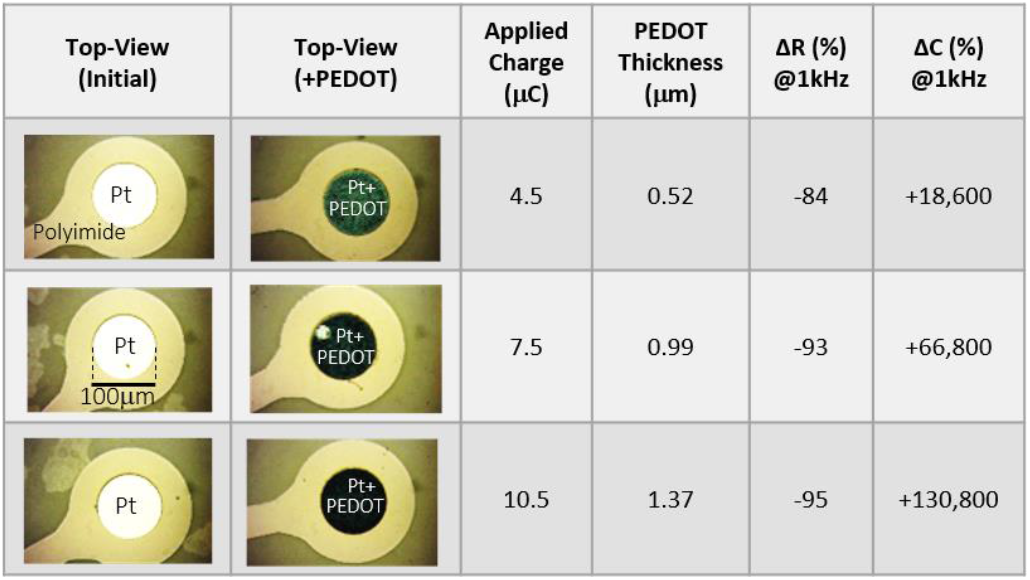
The coating properties that show how varying electroplating current densities result in various thicknesses for PEDOT films on platinum electrode sites alongside a close view of the electrodes before/after electroplating PEDOT.

| Top-View (Initial) | Top-View (+PEDOT) | Applied Charge ($\mu\text{C}$) | PEDOT Thickness ($\mu\text{m}$) | $\Delta R$ (%) @1kHz | $\Delta C$ (%) @1kHz |
| --- | --- | --- | --- | --- | --- |
|  |  | 4.5 | 0.52 | -84 | +18,600 |
|  |  | 7.5 | 0.99 | -93 | +66,800 |
|  |  | 10.5 | 1.37 | -95 | +130,800 |

**Table 2:**
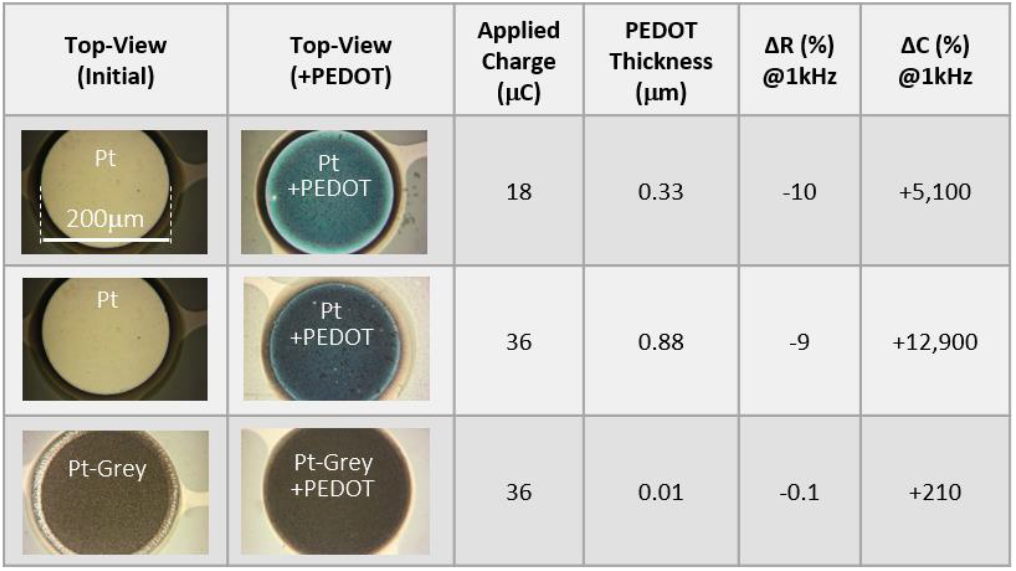
Coating properties and close view of the platinum electrode sites of the electrode arrays belonging to the Orion Cortical Prosthesis System made by SSMP and used in this work. The first two show PEDOT on Pt electrode sites and the third row shows PEDOT on Pt-Grey.

| Top-View (Initial) | Top-View (+PEDOT) | Applied Charge ($\mu\text{C}$) | PEDOT Thickness ($\mu\text{m}$) | $\Delta R$ (%) @1kHz | $\Delta C$ (%) @1kHz |
| --- | --- | --- | --- | --- | --- |
|  |  | 18 | 0.33 | -10 | +5,100 |
|  |  | 36 | 0.88 | -9 | +12,900 |
|  |  | 36 | 0.01 | -0.1 | +210 |

**Figure 3:**
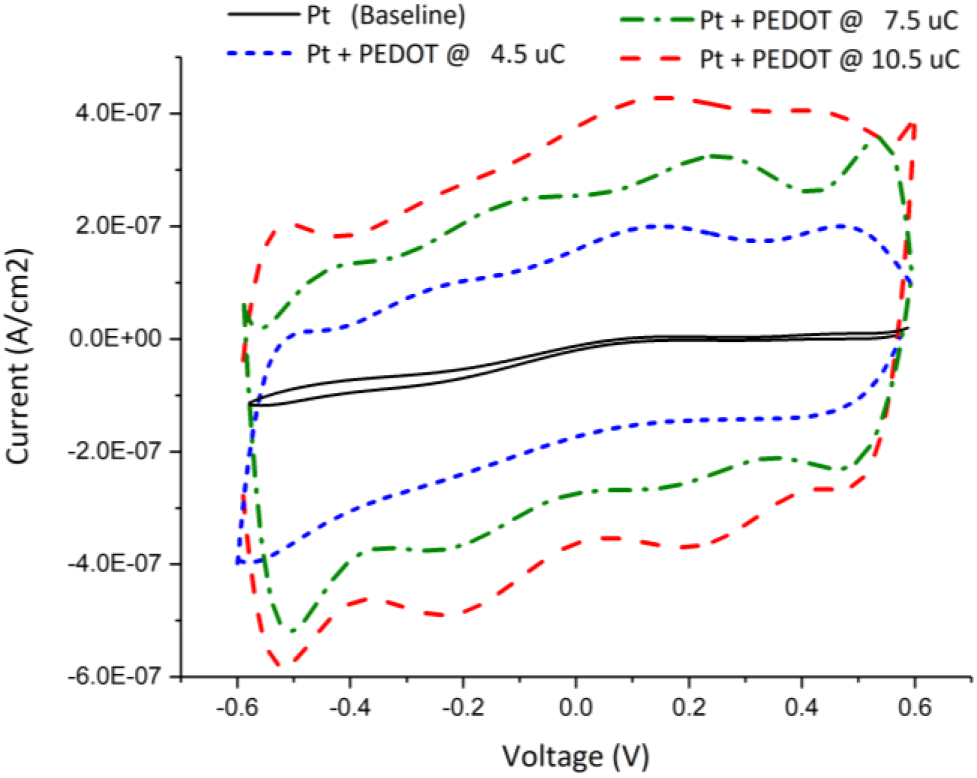
Averaged CV response of the platinum electrode sites before and after electroplating PEDOT thin films at 4.5μC, 7.5μC, and 10.5μC.

Fig. 4 depicts the CV results of the second experiment and compares the effects of coating PEDOT:BF4 on smooth platinum versus Pt-Grey. This comparison is carried out from the lens of the CV response at the electrode/electrolyte interface inside 10mM PBS. PEDOT alone on platinum improves the CSC (Fig. 4-blue) more than Pt-Grey alone does (Fig 3-green). Comparing the initial results, it was observed that PEDOT-on-Pt surpasses Pt-Grey-on-Pt by effecting an approx. 2.2x reduction in the interfacial electrical resistance and an approx. 3.7x increase in the interfacial electrical capacitance. These findings were key to the motivation behind this study which is to significantly improve upon the existing Pt-Grey coating technology developed by SSMP Inc. It was demonstrated that the combination of PEDOT on Pt-Grey exceeds either of the two individual coatings, offering superior enhancement of the desirable electrical properties while presenting a magnificent chemo-mechanical bond between the two. The results in Fig. 4 show that nearly 1μm of PEDOT on platinum benefits the electrochemical performance much better than 8 μm of Pt-Grey alone. PEDOT films thicker than 1μm do not proportionally increase the positive electrochemical performance impact.

**Figure 4:**
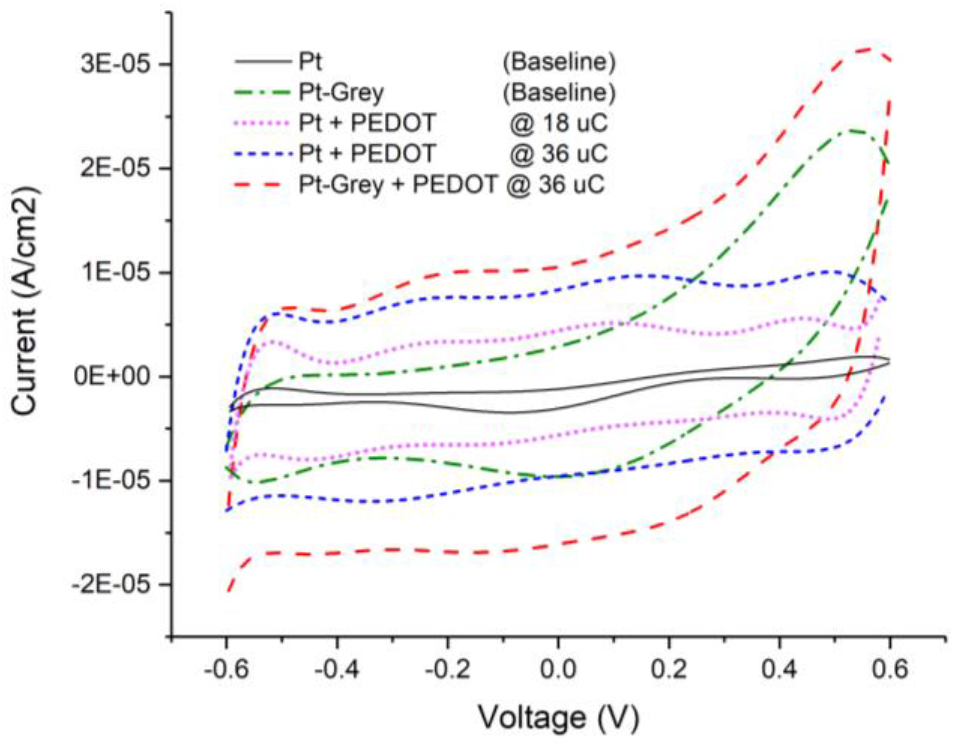
Averaged CV response of the platinum electrode sites with/without Pt- Grey before/after electroplating PEDOT thin films at 18μC and 36μC.

Indeed, the benefits of PEDOT coatings saturate at about 2μm. Other experiments showed that PEDOT coatings thicker than 2μm often suffer from mechanical instability and inconsistent quality.

Fig. 5 shows the *in vitro* long-term stability test-setup including the heating and electrical stimulation equipment. The results of that experiment are presented in Fig. 6. In all the cases, the impedance values are averaged from four similar electrodes (with diameters of 200μm) and yet separately electroplated and stimulated in PBS. In all the panels day-1 represents the blank electrodes while day-2 represents the electrodes after they are electroplated with their PEDOT coatings. The rest of the time the electrodes remain the same in terms of material configuration while they undergo continuous electrical stimulation. The results show that PEDOT coating on Pt electrodes decreases the initial resistive impedance by >3x and increases capacitance by more than 23x. Degradation in performance was observed starting from day 32 (Fig. 6-a&b). The Pt-Grey electrodes were also tested for long periods of time, but much longer than Pt electrodes. That’s because they are more robust and were similarly tested until degradation was observed. As shown in Fig. 6-c the presence of PEDOT decreases the electrical resistance by nearly 10% and diazonium salt helps decrease it even further, as expected. The enhancements remain intact until they start showing a decrease in their electrical performance at approximately 100 days. However they remain improved and functional for more than 450 days compared with no coating (blank Pt-Grey). Fig. 6-d shows the similar favourable impact of PEDOT and diazonium salt on Pt-Grey electrodes in terms of capacitive impedance. Again, after day-2 (applying the coatings) the impressive increase in capacitive impedance is observed for >3x with a robust endurance for up to 100 days before the degradation in the capacitive enhancement is observed. Though, the coatings still show some enhancement (~10% resistive and >2× capacitive) even after 450 days.

**Figure 5:**
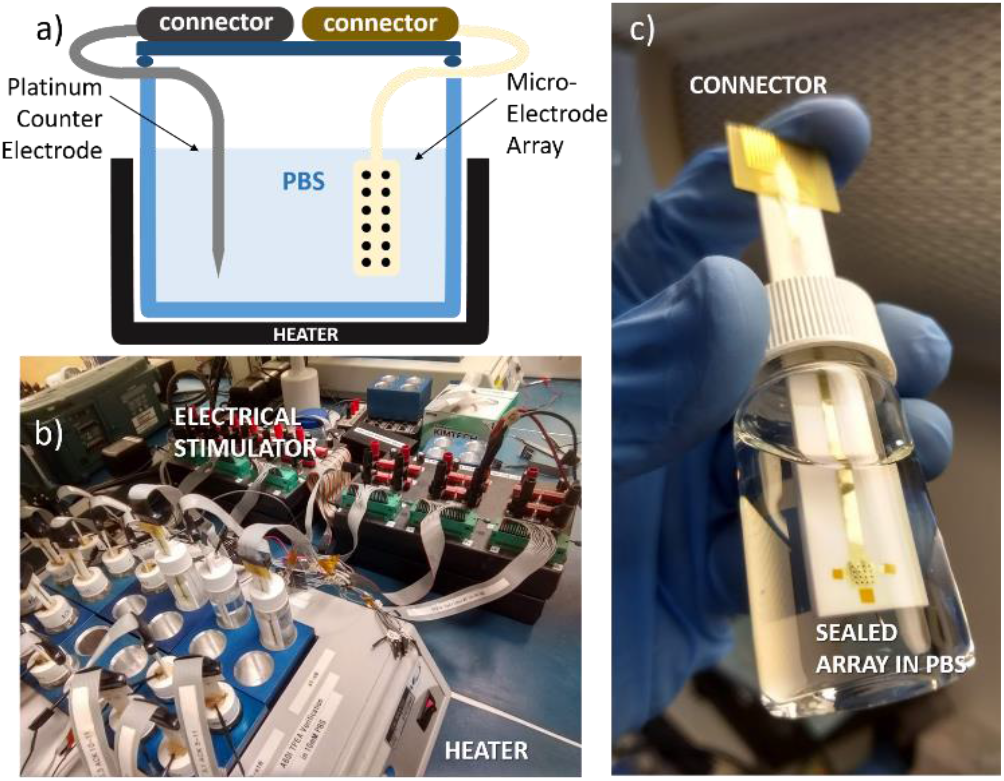
Long term *in vitro* stability test setup for the platinum electrode arrays, a) The schematic diagram shows the main components of the endurance test setup, b) The heating and stimulation units that provide electrical current and heat to the samples inside their PBS tubes while they are probed by oscilloscope for monitoring their biphasic waveforms, c) Close view of one of the samples inside its tube filled with 10mM PBS, sealed with silicone, and wired out for electrical connection.

**Figure 6:**
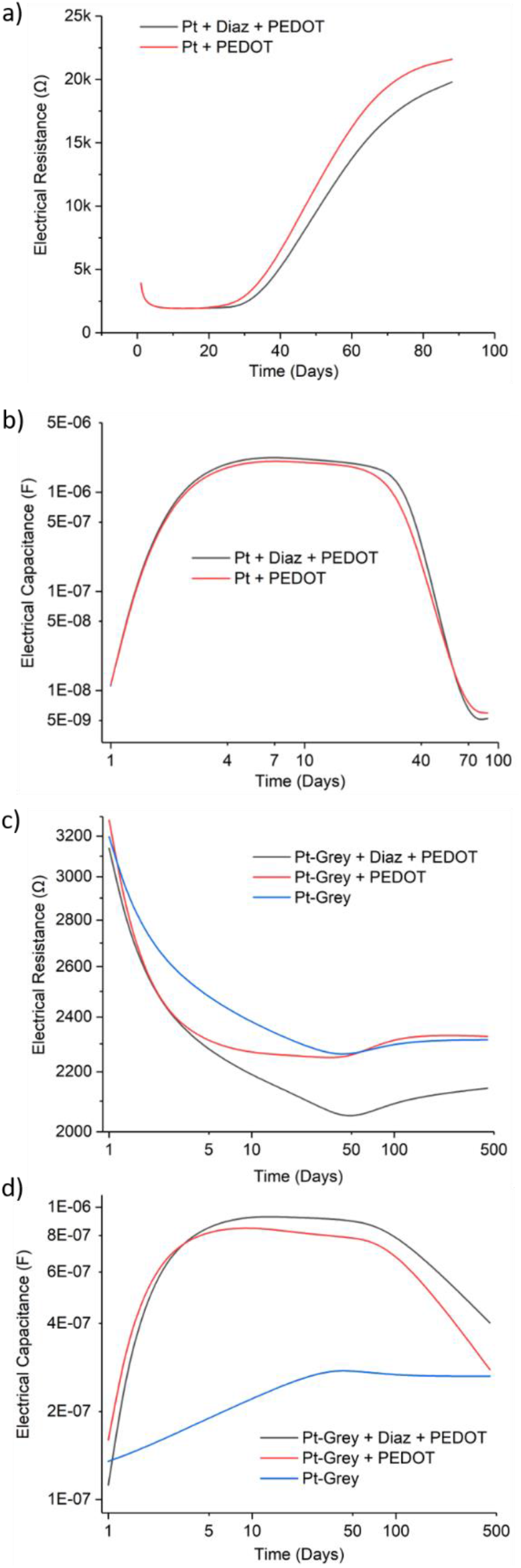
Long term *in vitro* stability test results at 37°C (accelerated) for the platinum electrode arrays with diameters of 200μm. a) 88 days of continuous electrical stimulation of Pt electrodes with PEDOT with/without diazonium salt and the comparison of its effect on electrical resistance profile over this period. b) 88 days of continuous electrical stimulation of Pt electrodes with PEDOT with/without diazonium salt and the comparison of its effect on electrical capacitance over this period. c) 452 days of continuous electrical stimulation of Pt-Grey electrodes with/without PEDOT and diazonium salt and its effect on electrical resistance over this period. d) 452 days of continuous electrical stimulation of Pt-Grey electrodes with/without PEDOT and diazonium salt and its effect on electrical capacitance over this period.

## DISCUSSION

Electrode arrays were designed, manufactured and provided by SSMP Inc. They are flexible and long-lasting electrodes that have been used in the Argus II Retinal Prosthetic Implant and implanted in the eyes of hundreds of human subjects for more than a decade, evoking action potentials that produce significant visual percepts [28]. With small variations in geometry, they have also been used in the Orion Cortical Prosthetic Implant introduced on the surface of visual cortices of six human patients delivering bionic vision [29]. The goal of this study was to enhance the electrical properties of SSMP’s existing implant electrode technologies. As expected, electroplating PEDOT:BF4 on platinum impressively enhanced the electrical characteristics of the electrode sites by decreasing the interfacial electrical resistance and increasing the electrical capacitance at the electrode/electrolyte interface. The endurance of Pt-Grey is much more satisfactory, especially at higher thicknesses which cannot be surpassed by the current state of the art of conductive polymers. Therefore, PEDOT alone cannot replace the benefits of Pt-Grey coatings entirely, but can be added to it. To take advantage of the impressive electrochemical and biological benefits of PEDOT and the outstanding electrochemical and mechanical endurance of Pt-Grey the two films were stacked and together with diazonium salt they created a hybrid coating that combined the best of both worlds. The complex intertwined microporosity of Pt-Grey entraps and houses the PEDOT molecules that are coated into its microporous geometry. This mechanical bond was strengthened by electrografting a chemical adhesion promoter called diazonium salt on to the Pt-Grey surface prior to the application of PEDOT. If the applied charge during electroplating of PEDOT on a given platinum electrode site with a diameter of 200μm is near the amount necessary to coat 1μm PEDOT, but instead applied on a site previously coated with 8μm Pt-Grey, no noticeable thickness is added to the overall coating composition. That is due to the integration of PEDOT molecules inside the vacancies of microporous geometry of Pt-Grey (See Fig.2-a). If the amount of charge being electroplated is more than this vacant volume of the given Pt-Grey film, the overall PEDOT coating starts to elevate above the surface of Pt-Grey and increases the overall thickness of the composite.

## MATERIALS AND METHODS

The electrode arrays that were used in this work are epiretinal and epicortical electrode arrays comprising 16 and 60-channel arrays with circular/planar sites in diameters of either 100μm or 200μm, respectively. They are both fabricated using single-layer sputtered platinum insulated by spin-coated polyimide that are patterned by standard photolithography using photoresist and ultra-violet light. As patented by SSMP Inc. the platinum electrode sites are coated with typically 8μm of Pt-Grey that enhances the microporosity, effective surface area and charge storage capacity above the minimum stimulation thresholds needed to evoke phosphenes and cause visual percepts. The specific settings for electroplating the Pt-Grey coating are intellectual property of SSMP Inc. and are not publicly available.

Electrografting diazonium salt on electrode surfaces was carried out based on a recipe originally reported by D. Chhin *et al.* [20]. Tetrafluoroboric acid, (4-thien-2-yl) aniline, diethyl ether, sodium nitrite, and lithium perchlorate were purchased from Milipore Sigma. Tetrafluoroboric acid 50% (124 μL, 1 mmol) was mixed with (4-thien-2-yl) aniline (175 mg, 1 mmol) with dropwise addition of acetonitrile until dissolution. The mix was then cooled to −10°C by placing the beaker inside a mixture of deionized ice-water and acetone. An aqueous solution of saturated sodium nitrite (69 mg, 1 mmol) was then added dropwise to the (4-thien2-yl) aniline solution. The resulting suspension was filtered using semi-permeable paper filter and rinsed with diethyl ether. Eventually, cyclic voltammetry was performed on microelectrodes that were placed inside a solution of 5 mM (4-thien-2-yl) diazonium salt dissolved in 100 mM LiClO4 and acetonitrile for 15 consecutive cycles sweeping from –0.5 V to 0.5 V at a step size of 0.1 V/s.

Electropolymerization of PEDOT was performed based on a previously published recipe by C. Bodart *et al.* [16]. Acetonitrile (ACN), polycarbonate (PC) (anhydrous, 99.7%), 3,4-Ethylenedioxythiophene (C6H6OS, 97%) and tetraethylammonium tetrafluoroborate (TEABF4, 99%) were purchased from Millipore Sigma. All chemicals were used as received. Electropolymerization was carried out galvanostatically in a 150mL of PC containing 426 mg EDOT monomer (20mM) and 3907 mg TEABF4 (120 mM). Prior to electropolymerization, all solutions were stirred and degassed for 20 minutes using nitrogen and a nitrogen blanket was maintained during the electropolymerization to limit the oxygen concentration in the solution and prevent unwanted oxidation. Electroplating was carried out in a three-electrode fashion using a Solartron Analytical 1287A potentiostat and galvanostat equipped with a 1252A Frequency Response Analyzer. Silver/silver chloride (Ag/AgCl) was used as the reference electrode, a platinum wire as the counter electrode and the platinum and Pt-Grey electrodes were connected as the working electrodes. The working electrodes were cleaned with 20% diluted sulphuric acid under cyclic voltammetry and then rinsed in deionized water. After electrodeposition, the electrodes were rinsed with deionized water, dried in air, and stored in ambient conditions.

The platinum electrodes with and without Pt-Grey were tested for long term durability under continuous round-the-clock electrical stimulation using charge-balanced, current-regulated, symmetric, biphasic pulse pairs at 120Hz and a peak current of 105μA. The long-term stability tests were carried out initially at 37°C in real time for 36 days and then 12.5 days at 67°C plus another 11 days at 87°C to accelerate the tests. According to the Arrhenius Equation [30] increasing the solution’s temperature by each 10°C accelerates the material aging by a power of two. That means the electrode materials at 67°C age 8x faster than when tested in a solution at 37°C. At 87°C the electrodes theoretically age 32x faster than 37°C. After the down conversions, 37°C equivalent total test durations of 88 days for Pt electrodes and 452 days for Pt-Grey electrodes were obtained.

## CONCLUSION

We introduced a novel thin film composition that takes advantage of two promising technologies and combines the best properties of both for enhancing flexible microelectrode arrays used in chronic neural stimulation. One is the most investigated and known conductive polymer, PEDOT, applied with an adhesion promoter called diazonium salt, and the other is Pt-Grey which is the key coating material used in the Argus and Orion visual prostheses. PEDOT not only enhances cellular agreement, but can promote cellular growth and carry various immunosuppressant drugs; it can also enhance the electrical properties of electrodes and surpass some of the best conventional inorganic coatings. However, PEDOT polymeric molecules do not stick well to their underlying substrates and this detracts immensely from their long-term durability. A few investigators have reported chemical approaches and partially improved PEDOT adhesion to some inorganic materials [17–19]. This report shows that if PEDOT is electroplated on microporous Pt-Grey, that was also previously treated with diazonium salt, it can endure round-the-clock continuous electrical stimulation in PBS at 37°C for more than 100 days without showing noticeable decline in the enhanced electrical properties (>2X better than Pt-Grey alone). These improvements are critical in designing the next generation of ultra-small and highly dense electrode arrays comprising hundreds or thousands of channels.

## ACKNOWLEDGMENT

The authors would like to acknowledge the financial support from the NSF supplemental funding under the award ECCS-1407880.

## DATA AVAILABILITY

The raw/processed data required to reproduce these findings cannot be shared at this time due to intellectrual property rights that belong to Second Sight Medical Products Inc.

## CONTACT

^*^M. H. M. Kouhani, tel: +1-517-7759527;

